# Identification and Validation of an inhibitor of the protein kinases PIM and DYRK

**DOI:** 10.1101/2025.10.22.683941

**Authors:** Gyula Bencze, Prabhadevi Venkataramani, Elad Elkayam, Keith D. Rivera, Ankur Garg, Istvan Szabadkai, Laszlo Orfi, Leemor Joshua-Tor, Darryl Pappin, Nicholas K. Tonks

## Abstract

Fermented wheat germ extract (FWGE), a nutraceutical with reported anticancer properties, contains numerous biologically active molecules, but its precise therapeutic constituents remain unclear. In this study, we identify and characterize a novel small-molecule inhibitor, CSH-4044, isolated from FWGE. Through preparative HPLC and structural elucidation via X-ray crystallography, CSH-4044 was revealed to be a unique benzothiazole compound. Kinase profiling demonstrated its high specificity toward the PIM and DYRK families of protein kinases. We determined the co-crystal structure of CSH-4044 bound to PIM1, revealing ATP-competitive binding, and critical hydrophobic and hydrogen-bonding interactions. A chemically synthesized version of CSH-4044 mirrored the activity of the natural product, confirming structural integrity and biological equivalence. Functionally, CSH-4044 suppressed PIM3-driven BAD phosphorylation in pancreatic cancer cells and reduced DYRK1A-mediated Tau phosphorylation in neuronal cells. Our findings position CSH-4044 as a promising lead for targeting PIM and DYRK families of kinases and highlight FWGE as a source of potential therapeutic compounds.

## INTRODUCTION

In recent years, nutraceuticals are gaining importance as therapeutic agents for cancer due to their low toxicity and promising effects in patients^1^. One such nutraceutical, a natural derivative of fermented wheat-germ extract (FWGE), has been used to treat cancer patients undergoing chemotherapy or radiotherapy in Eastern European countries^2^. It has been reported to be effective against different cancer types, including lung, colon, prostate, and breast cancer^3^. The anti-proliferative activity of FWGE has also been studied in various animal models and cancer cell-lines and is shown to trigger tumor cell-death in a dose-dependent manner^4,5^. One such study demonstrated that FWGE in combination with cisplatin had antiproliferative effects and potentiated cisplatin-induced apoptosis in ovarian cancer^4^. In addition, some studies have also investigated its potential to inhibit cell migration and invasion in oral squamous cell carcinoma^5^.

The production process of FWGE involves fermentation of wheat germ by *Saccharomyces cerevisiae* followed by liquid separation, drying, and granulation^6^. Like other nutraceuticals, FWGE likely contains a wide array of different molecules. Thus, exploiting its full potential will require a more precise definition of the active core components of the mixture and the biochemical characterization of their mechanism of action. Recent studies indicate that two quinones, 2-methoxy benzoquinone and 2,6-dimethoxy benzoquinone, might contribute to the biological activity of FWGE^7^. In this study, in which we sought to identify additional active components of the extract, we generated different fractions of FWGE using preparative HPLC. Using subfraction A250, we have purified and crystallized CSH-4044, a novel protein kinase inhibitor.

In this study, we identify and characterize CSH-4044, a unique nutraceutical-derived small molecule that functions as an ATP-competitive inhibitor of PIMs and DYRKs^8,9^. We solved its structure by X-ray crystallography, established its mode of binding to PIM1, and confirmed its activity through both natural product isolation and synthetic reproduction. Functionally, we demonstrate that CSH-4044 reduces phosphorylation of BAD in pancreatic cancer cells and decreases Tau phosphorylation in neuronal cells, thereby validating its kinase inhibitory activity in two disease models. Our findings highlight FWGE as a source of structurally novel bioactive compounds and establish CSH-4044 as a promising lead scaffold for therapeutic development against PIM- and DYRK-driven pathologies.

## RESULTS

### Enrichment of active fractions from fermented wheat germ extract (FWGE)

In a previous study from our lab, we isolated Fraction A250 through bioassay-guided fractionation and alcohol extraction from fermented wheat germ extract (FWGE). Although A250 constituted only 3% of FWGE, it showed antiproliferative activity against different cancer cell lines^10^. Liquid–liquid extraction of Fraction A250 with ethyl acetate yielded two distinct subfractions: a hydrophilic aqueous phase (A251) and a lipophilic organic phase (A252). HPLC confirmed that flavonoid glycosides partitioned into A251, whereas dimethoxy benzoquinone (DMBQ) and other components were enriched in A252 (Fig S1A-B). Bioactivity testing showed that A252 exhibited greater cytotoxic potential than either A250 or A251 (Fig S1C), suggesting enrichment of active compounds in the lipophilic fraction. To determine the role of DMBQ, Fraction A252 was treated with ascorbic acid (ASC) in methanol to reduce DMBQ to 2,6-dimethoxyhydroquinone (DMHQ), allowing separation by HPLC (Fig S1D). Although the DMBQ-depleted material (Fraction A252-EP) showed reduced in vitro activity compared with A252, the persistent cytotoxicity of this still-complex fraction suggested the presence of additional, unidentified bioactive components.

### Kinase Selectivity Profiling of A250-Derived Fractions

As part of the optimized extraction and purification workflow (Fig 1A), we concentrated the small-molecule content of FWGE into Fraction A250. Despite its low abundance, Fraction A250 retained nearly all the cytotoxic activity observed in crude FWGE. To assess their potential to inhibit protein kinase targets, Fraction A250 and its subfractions, including the hydrophilic A251 and the lipophilic A252, were screened at 1µg/ml concentration using a 138-kinase panel (The International Centre for Kinase Profiling, University of Dundee). Initial analysis revealed that whereas A250 exhibited inhibitory activity against several kinases, most of the inhibitory effects localized to Fraction A252, the ethyl acetate-soluble fraction enriched in lipophilic small molecules.

**Figure 1.**
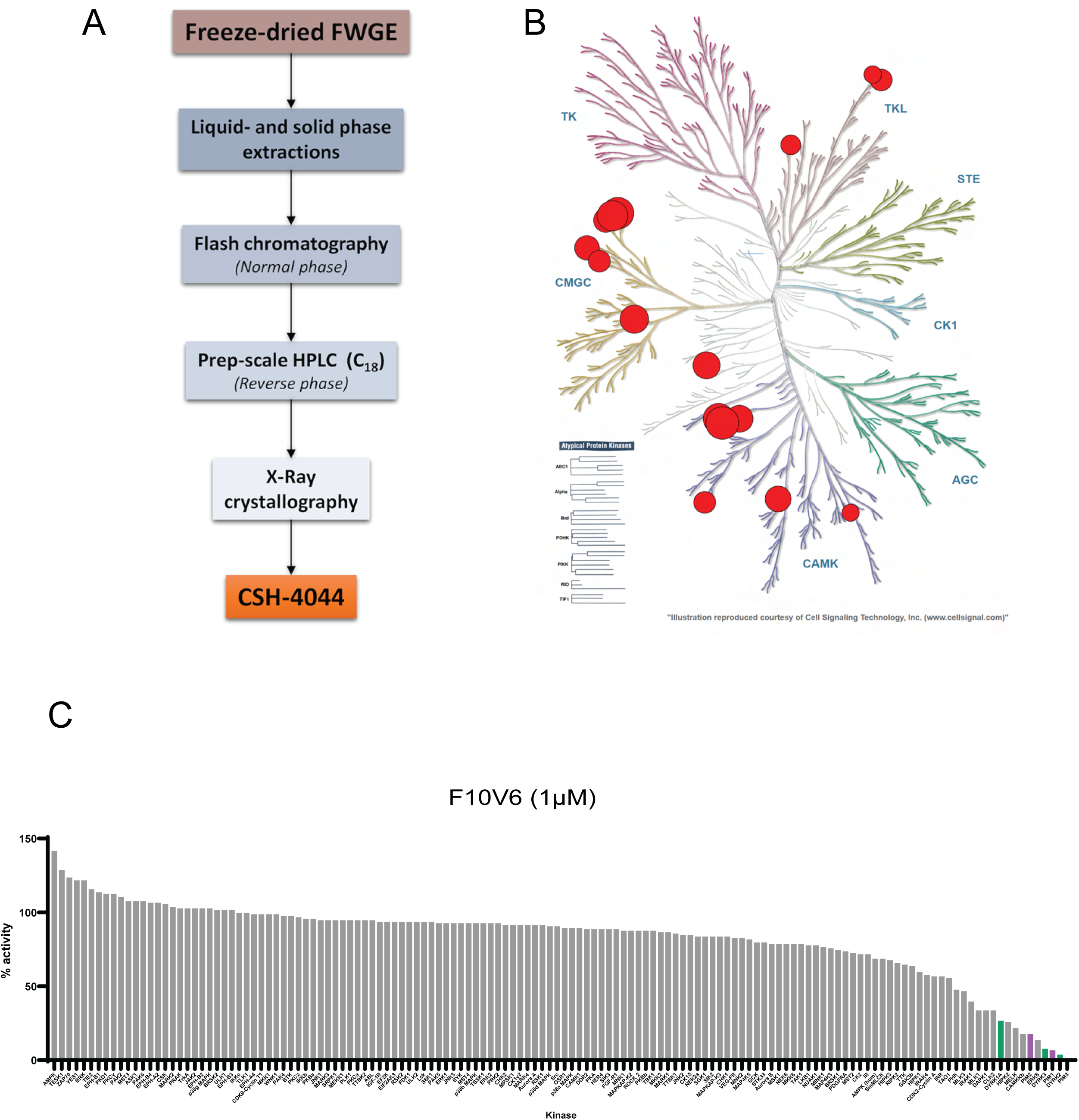
Kinase Selectivity Profiling of A250-Derived Fractions. **(A)** Extraction and purification workflow for the isolation of CSH-4044 from Freeze-dried FWGE. **(B)** Kinome tree visualization of the inhibitory effect of CSH-4044 across a panel of about 140 kinases. There was no significant effect on 130 of the kinases that were tested. At a concentration of 1µM, PIM3 was inhibited by 99%, PIM1, DYRK2 and DYRK3 by ∼95% (large circles). Inhibition of the related enzymes, ERK8, CAMKK2, MELK and HIPK2 was detected, but to a lesser extent (small circles) **(C)** Waterfall plot of the kinase selectivity panel results for the isolated compound F10V6 from FWGE (1µM). The PIM and DYRK kinases are labelled in purple and green respectively.

Subsequent fractionation and kinase profiling of A252-derived subfractions led to the identification of a compound with a highly selective inhibitory profile, primarily targeting the PIM and DYRK families of kinase (Table 1, Fig 1B). The FWGE-derived compound exhibited markedly improved selectivity (Fig 1C). These findings suggest that the biologically active compound in FWGE may represent a novel and selective chemical scaffold for kinase inhibition.

**Table 1.**
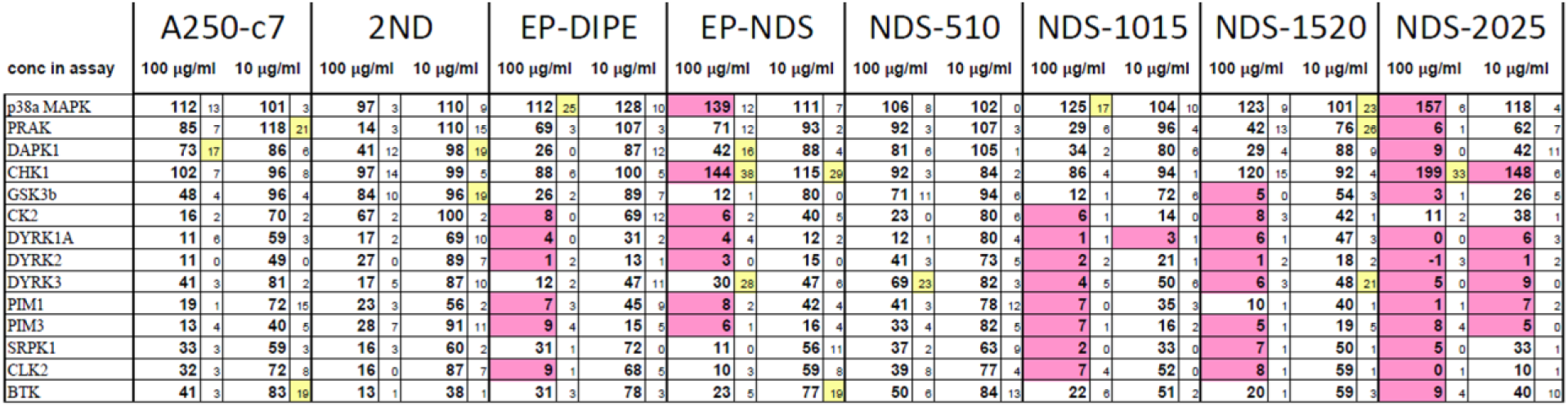
Inhibitory activity of different subfractions of FWGE against protein kinases. The subfractions containing CSH-4044 primarily targeted PIMs and DYRKs.

### Preparative Isolation of Active Subfractions

Due to the pronounced kinase-inhibitory profile of Fraction A252 in a 138-kinase panel screen, especially against PIM3 and DYRK2, larger quantities were processed for milligram-scale isolation of individual compounds. A combination of preparative-scale, normal-phase flash chromatography, reverse-phase HPLC, and MS-coupled analytical HPLC was employed to isolate purified components from freeze-dried FWGE, yielding Subfractions 2025A, B, and C (Fig 2A-B). All three subfractions displayed measurable kinase inhibition, with 2025A producing the lowest IC₅₀ values against both PIM3 and DYRK2 (Fig 2C). Subsequent HPLC analysis of 2025A under alternative conditions revealed two distinct peaks. Further separation yielded two purified compounds, one of which, designated F10V6, showed markedly higher kinase-inhibitory activity (Fig 2D).

**Figure 2.**
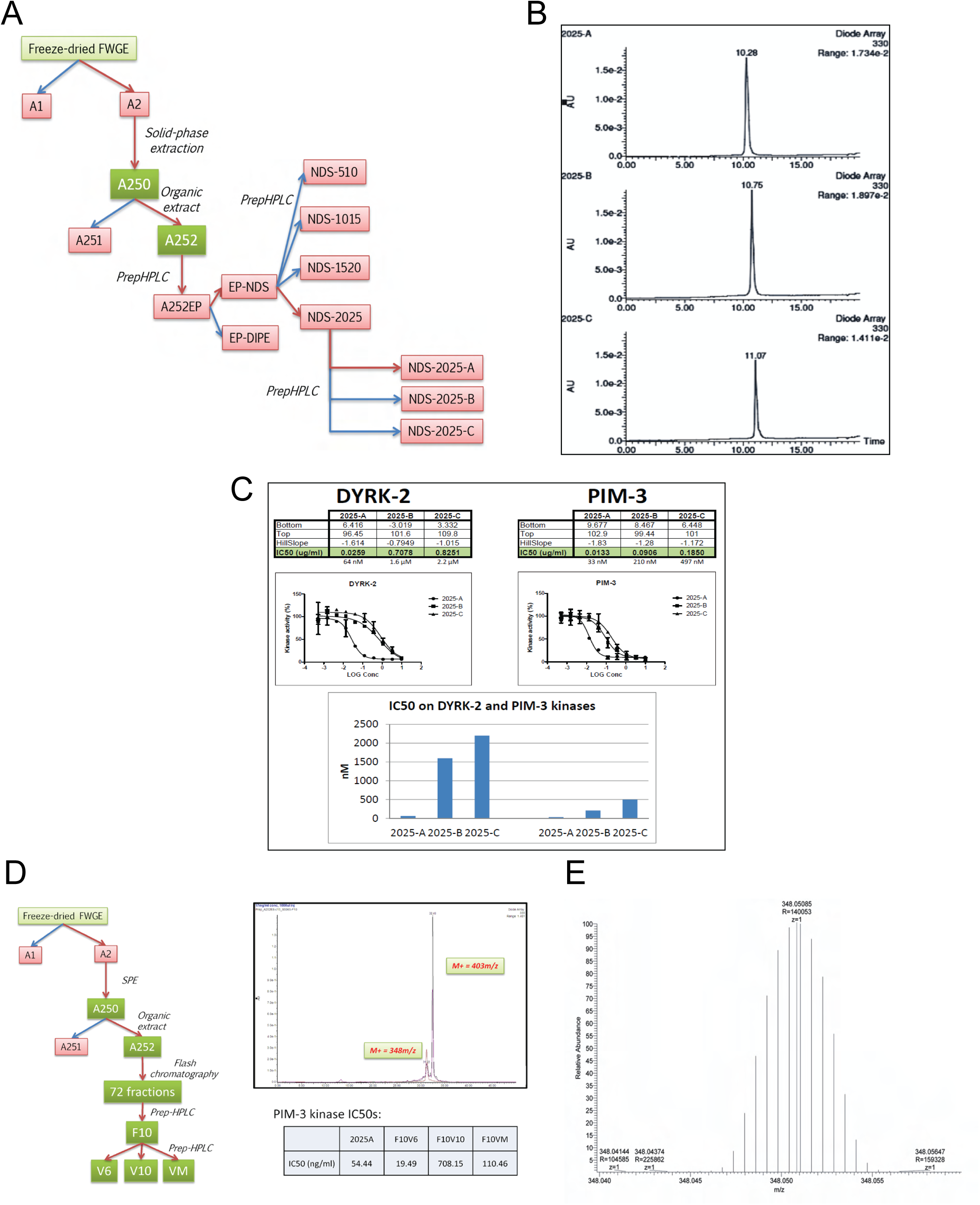
Preparative isolation of the different fractions of A250. **(A)** Flowchart outlining the steps involved in the isolation of fractions from FWGE with the highest activity **(B)** HPLC-UV chromatogram of the fractions – 2025-A, 2025-B and 2025-C. **(C)** Inhibitory effect of 2025-A, B and C on PIM and DYRK kinases **(D)** Flowchart outlining the isolation of F10V6 from FWGE, its molecular weight and IC50 against PIM3 **(E)** Orbitrap Q-TOF data of F10V6

To determine the identity of F10V6, the molecular weight was first obtained via electron ionization mass spectrometry (EI-MS) (Fig 5B). Fragmentation patterns were then analyzed using a triple-quadrupole MS, and high-resolution mass data were collected via Orbitrap Q-TOF, with mass accuracy to four decimal places (Fig 2E, Fig S1E-F). Searches against the NIST compound database revealed no matches with >33% probability, suggesting that F10V6 may represent a novel small molecule.

### F10V6 is a benzothiazole with a unique molecular structure

To determine the structure of the active component isolated from FWGE fraction A252, the extract was crystallized at room temperature, which generated orange crystals (Fig 3A). X-ray crystallography analysis revealed that the compound was a benzothiazole, 5-Dihydroxy-4-methoxy-phenyl)-(6-hydroxy-5-methoxy-benzothiazol-2-yl)-methanone with a molecular weight of 347.3415 (Fig 3B). Although a ChemSpider search for the molecular formula of the compound (C_16_H_13_NO_6_S) showed more than 2880 structures that shared the same molecular weight, and 195 structures with the same molecular formula, F10V6 turned out to be a unique benzothiazole, since none of the hits generated could match its structure. After solving the structure of the isolated molecule, we assigned the name CSH-4044.

**Figure 3.**
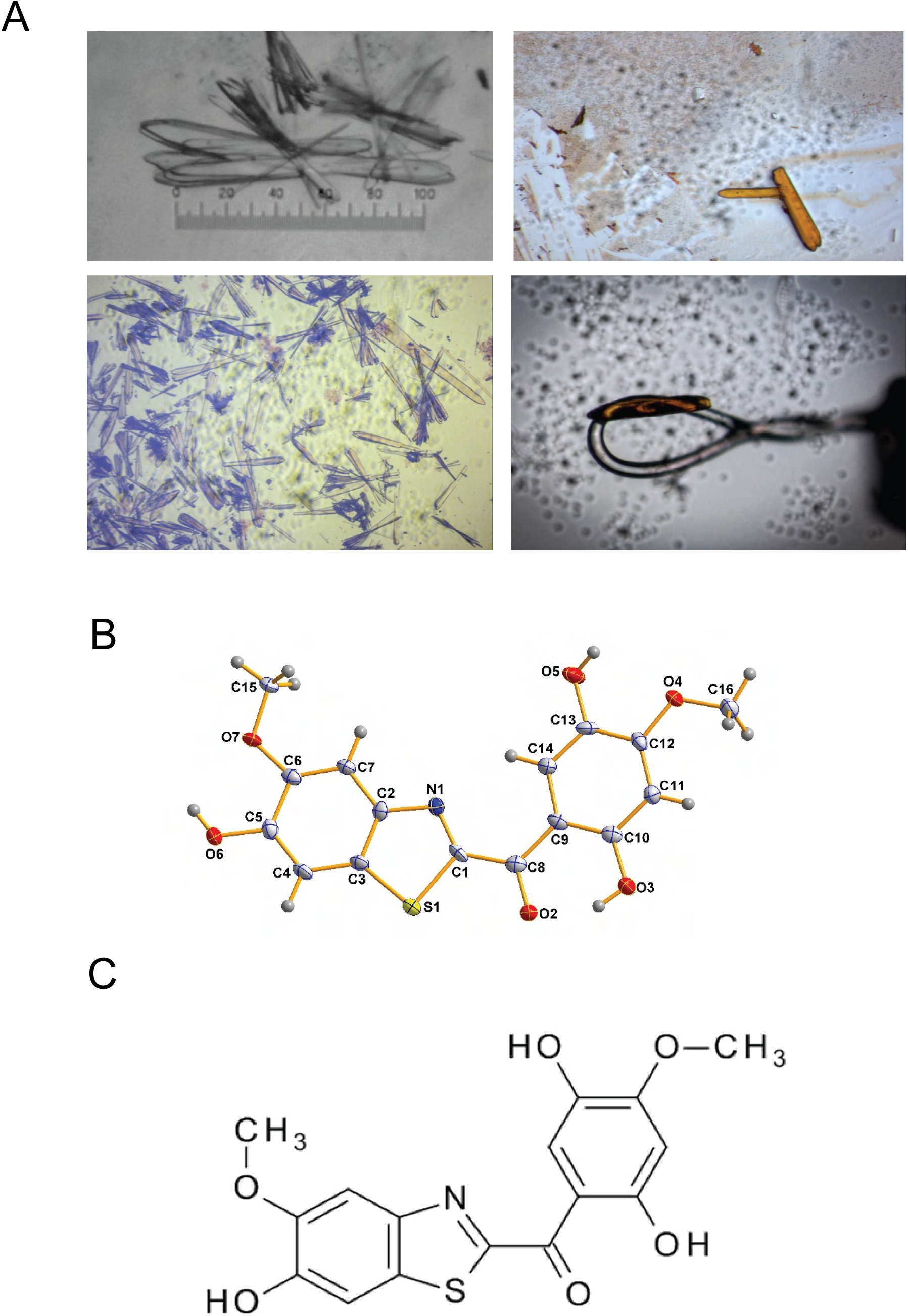
F10V6 is a benzothiazole with a unique molecular structure. **(A)** Microscopic view of the generated orange crystals of F10V6 **(B)** Molecular structure of F10V06 ((2,5-Dihydroxy-4-methoxy-phenyl)-(6-hydroxy-5-methoxy-benzothiazol-2-yl)-methanone

### Crystal structure of PIM1-CSH-4044 complex

To investigate the structural basis of CSH-4044-mediated PIM1 inhibition, we determined the crystal structure of PIM1 in complex with CSH-4044. The PIM1-CSH4044 cocrystals belong to space group P6_5_ with one complex in the asymmetric unit. PIM1 has a typical bi-lobal serine/threonine kinase fold (Fig 4A, Table 2). A clear difference in electron density for CSH-4044 was observed, allowing us to place the inhibitor unambiguously in the ATP-binding pocket in a flat orientation between the two lobes. The nucleotide binding pocket is lined by several hydrophobic residues (Leu44, Val52, Phe49, Ile104, Val126, Leu174, Ile185) that surround the inhibitor. Moreover, the benzothiazole 5-methoxy group of CSH-4044 forms an H-bond with Lys67, whereas its 6-hydroxyl group H-bonds with Lys67 and the main-chain amide of Asp186. Additionally, the 6-hydroxyl group is stabilized by water-mediated interactions with Glu89 and the main-chain amide of Phe187 (Fig 4B). Similarly, the 2-hydroxyl group on the phenyl ring of CSH-4044 establishes water-mediated interactions with the Arg122 and Pro123 carbonyls, whereas its 4-methoxy group undergoes water-mediated interactions with Asp131 and Asp128 amide groups (Fig S2A). Notably, these hydrophobic interactions are very similar to the interaction with the adenine base in the PIM1-AMP-PNP structure (PDB 1XR)^11^ (Fig S2B), with the exception of P-loop Phe49, which repositions to occupy the b-phosphate position of AMP-PNP and closes the nucleotide binding pocket, as observed in the PIM1-Staurosporine structure (PDB 1YHS)^12^ (Fig S2C). However, due to the polar nature of the CSH-4044, the direct H-bond interactions with Lys67 and Asp186 main-chain amide appear to be CSH-4044-specific. Additionally, a Mg^2+^ ion is required for catalysis by PIM1 and is coordinated in an octahedral geometry via direct interactions with Ser189 and 5 water molecules, whereas the catalytic Asp198 interacts directly with the water molecules coordinating the Mg^2+^ octahedron (Fig 4C).

**Figure 4.**
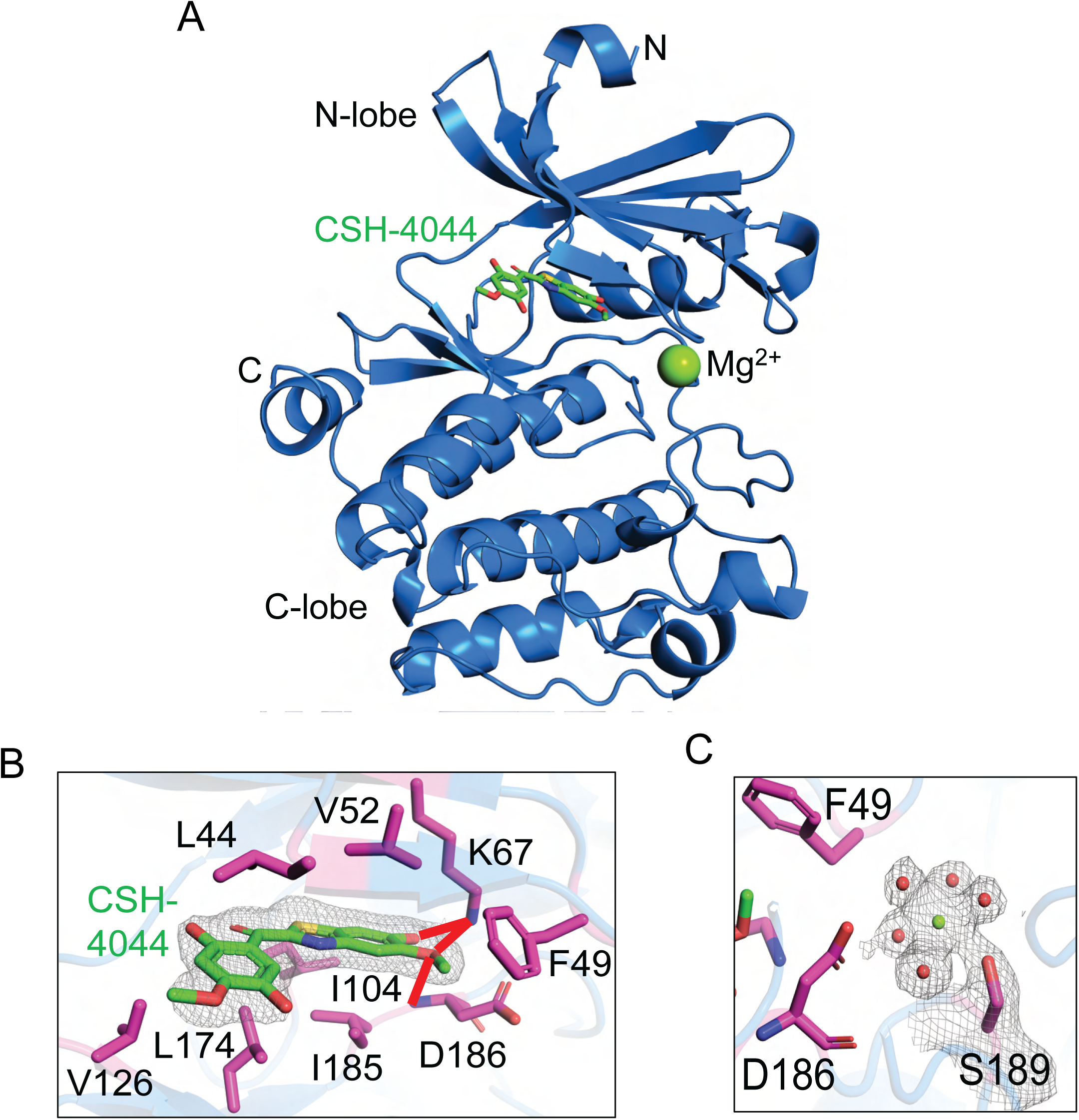
Crystal structure of human PIM1-CSH4044 complex. **(A)** A cartoon representation of the PIM1 crystal structure in complex with the small molecule inhibitor CSH-4044, shown as sticks (green). **(B)** A close-up view of CSH-4044 in the PIM1 nucleotide-binding site. The inhibitor is supported by several hydrophobic residues and direct H-bonds (red dotted lines). **(C)** The observed 2Fo-Fc electron density for the coordinated Mg^2+^ ion (green sphere) in the PIM1-CSH-4044 complex is shown as mesh at 1.0 s level (carve radius of 1.6 Å) along with coordinating water molecules (red spheres). Ser189 directly interacts with the Mg2+ ion, while Asp186 coordinates surrounding water molecules.

**Table 2.** X-ray data collection and structure refinement statistics for the crystal structure of the PIM1-CSH-4044 complex.

|  | PIM-CSH-4044 |
| --- | --- |
| Beamline | 8.2.1 |
| Wavelength (Å) | 1.0 |
| Data collection on | 051815 |
| Space group | P6<sub>5</sub> |
| a, b, c (Å) | 97.93, 97.93, 80.40 |
| $\alpha$, $\beta$, $\gamma$ (°) | 90, 90, 120 |
| Resolution (Å) | 41.82-1.94<br>(2.01-1.94) |
| No of reflections | 1,70,776 |
| Unique reflections | 32,068 |
| R<sub>merge</sub> (%) | 4.7 (56.3) |
| $\langle I/\sigma(I) \rangle$ | 20.6 (2.8) |
| CC<sub>1/2</sub> | 0.99 (0.81) |
| Completeness (%) | 99.77 (99.94) |
| Multiplicity | 5.3 |
| Refinement |  |
| R<sub>work</sub>/R<sub>free</sub> | 0.161/0.192 |
| Protein | 274 |
| Nucleic acid | 0 |
| Hetero atoms | 3 |
| Solvent atoms | 177 |
| Bond lengths (Å) | 0.018 |
| Bond angles (°) | 1.55 |
| Most favored (%) | 98.14 |
| Allowed (%) | 1.49 |
| Outliers (%) | 0.37 |
| Rotamer outlier (%) | 0.4 |

<sup>a</sup>Values in parentheses represent the highest resolution shell.

### Chemical Synthesis, Structural Verification, and quantification of CSH-4044

A core structural search using SciFinder revealed that although over 230 structural analogs of the lead inhibitor had been reported, the specific compound was not commercially available. To enable further functional studies, the compound was chemically synthesized (Vichem Chemie Ltd., Budapest, Hungary) (Fig 5A). The synthetic product was purified to >99% using a Waters preparative HPLC system. Its structure was confirmed by NMR spectroscopy and mass spectrometry, matching the spectral features of the natural product isolated from FWGE (Fig S3A-B).

**Figure 5.**
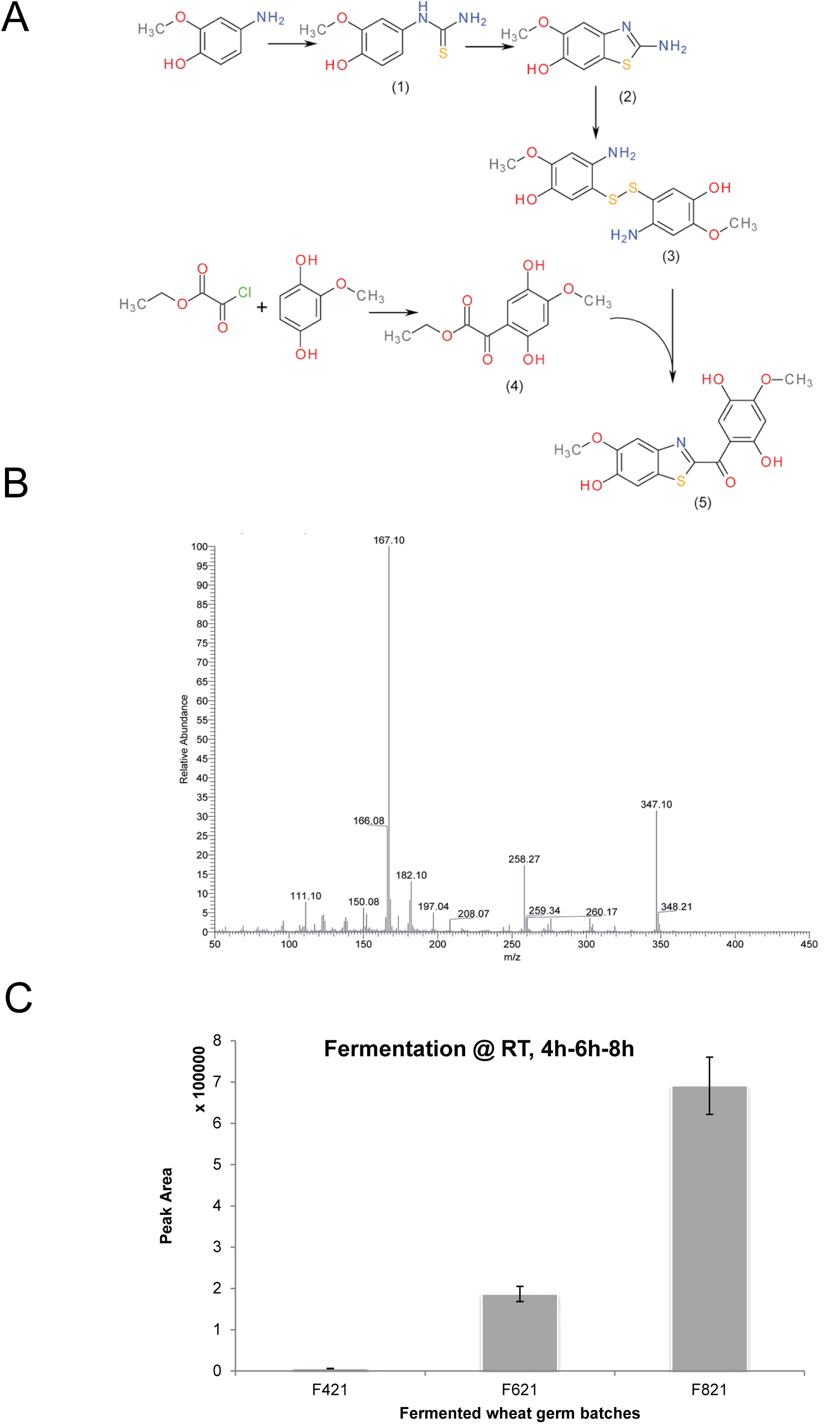
Chemical Synthesis, Structural verification and quantification of CSH-4044. **(A)** Flowchart outlining the steps involved in the chemical synthesis of CSH-4044. **(B)** Electron ionization mass spectrum of the isolated compound, showing the major fragment ions. **(C)** HPLC-MS analysis showing the abundance of CSH-4044 during the fermentation process at room temperature in three different time points.

To validate the synthetic route functionally, the purified compound was tested alongside the natural isolate in kinase inhibition assays. The synthetic version (CSH-4044) retained the same level of biological activity as the naturally derived F10V6, with comparable IC₅₀ values against PIM3 and DYRK2 (Table 3). These results confirmed that the synthetic compound replicates the in vitro kinase inhibitory activity of its natural counterpart.

**Table 3.**
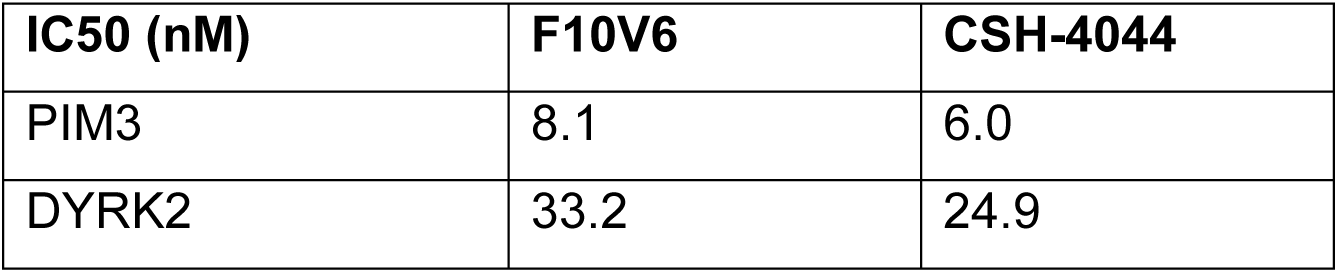
Comparison of inhibitory activity of F10V6 isolated from FWGE with synthetic CSH-4044 against PIM and DYRK kinases.

| **IC50 (nM)** | **F10V6** | **CSH-4044** |
| --- | --- | --- |
| PIM3 | 8.1 | 6.0 |
| DYRK2 | 33.2 | 24.9 |

To assess the production dynamics of CSH-4044 during FWGE fermentation, its concentration was quantified in the broth using HPLC-MS peak area analysis (Fig 5B). The compound’s abundance increased over time, with a higher signal detected at 8 hours compared to 4 hours of fermentation (Fig 5C) indicating that the fermentation step was crucial for the compound synthesis from FWGE.

### CSH-4044 inhibits PIMs and DYRKs in two cell-based models

To confirm the intracellular relevance of the identified kinase inhibitor, multiple human cell lines were tested for endogenous expression of PIMs (PIM1, PIM2, PIM3) and their downstream substrate, the pro-apoptotic protein BAD^13–16^. Among the eight lines tested, MiaPaCa-2, a human pancreatic cancer cell line, exhibited significant levels of all four proteins (Fig S4), making it suitable for functional validation. PIMs have been shown to inhibit apoptosis by phosphorylating pBAD at Ser112 in pancreatic cancer. Phosphorylation of BAD leads to its impaired binding to Bcl-XL and Bcl-2. The presence of unbound Bcl-XL eventually leads to inhibition of apoptosis. Elevated PIM3 levels have also been shown to increase BAD Ser112 phosphorylation, leading to apoptosis inhibition^13,14,17^. Treatment of MiaPaCa-2 cells with CSH-4044 markedly reduced phosphorylation of pBAD at Ser112 in MiaPaCa-2 (Fig 6A, 6B).

**Figure 6.**
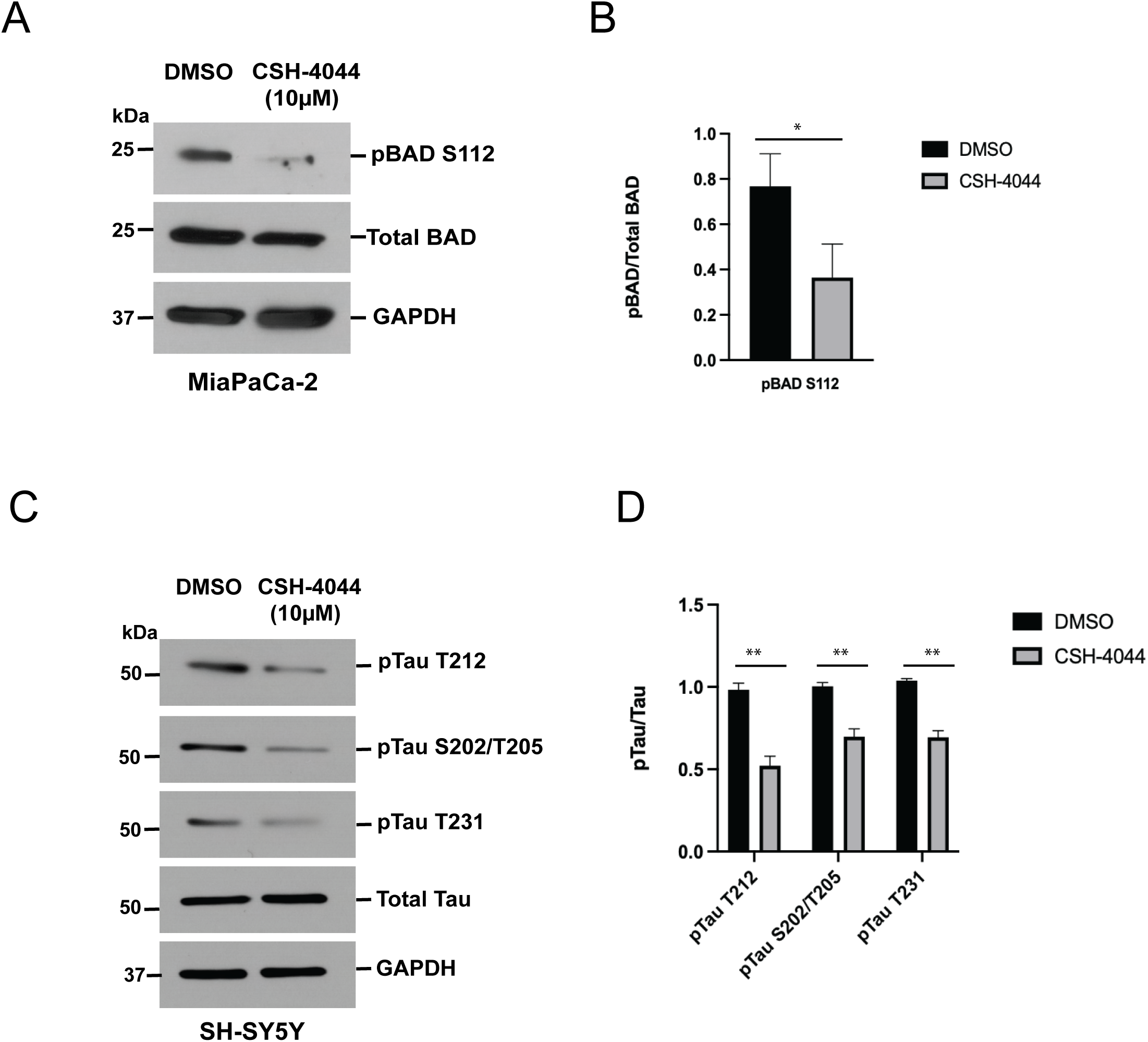
CSH-4044 inhibits PIMs in a Pancreatic Cancer Model. **(A-B)** CSH-4044 (10µM) reduces phosphorylation of BAD Ser112 in MiaPaCa-2 compared to the DMSO control. **(C-D)** CSH-4044 markedly reduces Tau phosphorylation at Ser212, Ser202/205 and Thr231 in SH-SY5Y cells (P value – 0.0012).

Since, CSH-4044 is a dual PIM and DYRK kinase inhibitor, we also tested whether CSH-4044 can affect the phosphorylation of Tau at multiple residues. DYRK1A has been shown to hyperphosphorylate Tau leading to the formation of neurofibrillary tangles in Alzheimer’s disease. The ability of Tau to promote microtubule assembly was markedly reduced due to its phosphorylation by DYRK1A^18–20^. Using the cell-line SH-SY5Y, we observed that CSH-4044 could reduce Tau phosphorylation at three residues, Tau S212, S202/T205 and T231 (Fig 6C, 6D). This indicates that CSH-4044 could target PIMs and DYRKs in two cell-based models.

## DISCUSSION

FWGE is a promising therapeutic agent for cancer and has been shown to exert anti-proliferative and cytotoxic effects in several cell-based models^21–23^. In this study, we successfully purified and identified a small-molecule inhibitor named CSH-4044 from a fraction A252 of FWGE and characterized it as a specific inhibitor of the PIM and DYRK families of kinases using peptide-based kinase activity assays.

Considering the significance of PIMs in cancer, we determined the crystal structure of PIM1 in complex with CSH-4044. Several independent groups have reported the crystal structure of PIM1 and PIM2 kinases either in the presence or absence of inhibitors^24–26^. In agreement with previous crystal structures, we report that the PIM1 structure assumes a two-lobe kinase fold with a deep intervening cleft. The two lobes are connected by the hinge region, wherein several residues have been identified to bind ATP^27^. The ATP-binding pocket of PIMs adopts an open conformation, indicating that they are constitutively active^25,26^. PIMs are structurally unique due to the presence of a proline residue (Pro123) in the hinge region of their ATP-binding site^28,29^. The proline residue hinders the formation of a second bond between PIMs and ATP, which is not usually observed in the case of other kinases^11,26^. Our structural studies reveal that CSH-4044 is an ATP-competitive inhibitor of PIM1 kinase. This suggests that CSH-4044 might be valuable in exploiting the novel ATP-binding site of PIMs to optimize and develop specific small-molecule inhibitors.

Most alterations in cancer are associated with signaling pathways that control cell growth and division, cell motility, cell death, and cell fate. Their dysregulation is often responsible for mediating distortions of wider signaling networks fueling cancer progression ^30^. Kinases often form part of the commonly altered signaling pathways in cancer ^31^. The PIM family of kinases has been implicated in tumorigenesis due to their constitutive activity and ability to regulate multiple survival and growth pathways^24,32–40^. PIM3 contributes to pancreatic ductal adenocarcinoma progression by phosphorylating the pro-apoptotic protein BAD at Ser112, thereby blocking its apoptotic activity^17,41,42^. We show that CSH-4044 inhibits PIM activity *in vitro* and reduces phosphorylation of BAD in MiaPaCa-2, supporting a direct intracellular effect.

In addition to PIMs, CSH-4044 demonstrated selective inhibition of the DYRK family of kinases, including DYRK1A. DYRK1A phosphorylates a range of substrates^9,43–46^, including Tau, where it promotes pathological hyperphosphorylation linked to Alzheimer’s disease^18–20^. In SH-SY5Y cells, treatment with CSH-4044 decreased phosphorylation of Tau at multiple residues, consistent with the inhibition of DYRK activity. These results extend the biological activity of CSH-4044 beyond oncology and highlight its potential utility in modulating kinase pathways associated with neurodegeneration.

Taken together, this study establishes CSH-4044 as a unique benzothiazole scaffold with dual inhibitory activity against PIM and DYRK kinases. Our results also highlight FWGE as a source of structurally novel bioactive compounds and underscore the potential of nutraceutical-derived metabolites to yield selective small-molecule kinase inhibitors.

## CONCLUSION

Despite the importance of PIM as a target in cancer and the fact that ATP-competitive, small molecule PIM inhibitors have been tested in clinical trials against different cancers, including prostate, myeloma, lymphoma, and acute myelogenous leukemia, to date, there are no FDA-approved PIM inhibitors^28,29,47^. Consequently, the development of PIM inhibitors is an unmet need. Our study identifies CSH-4044 as a novel, dual-specific ATP-competitive inhibitor of PIMs and DYRKs, with therapeutic potential in pancreatic cancer and neurodegenerative diseases such as Alzheimer’s. These findings underscore the value of nutraceutical-derived scaffolds as a source of structurally unique, biologically potent drug candidates.

## MATERIALS & METHODS

### FWGE Component Separation and Extraction

Commercially available freeze-dried fermented wheat germ extract (FWGE) (10 gms) obtained from American Biosciences, Inc., was resuspended in 30mL analytical grade methanol (Sigma-Aldrich) and sonicated for 5 min. The insoluble solids were separated with a glass Buchner filtering funnel combined with a quantitative grade filter paper. The pellet was resuspended (Fraction A1) in methanol and filtered again. The filtrates were combined, and a Büchi Rotavapor R-100 bench-top evaporator system was used to remove the solvent, leaving the soluble part of FWGE behind. (Fraction A2). Fraction A2 was dissolved in 80 mL of distilled water and was passed through a solid phase extraction (SPE) column (Hydrophile Lipophile Balance, HLB, Waters). The column was washed with 50 mL of water, and the isolated components from the column were eluted with 50% methanol. The excess methanol was removed with the Büchi Rotavapor R-100 bench-top evaporator system. The samples were flash frozen in liquid nitrogen and lyophilized using Labconco FreeZone 6 Plus freeze-dryer (Fraction A250). Next, the fraction A250 was resuspended in 200 mL of distilled water, and its pH was adjusted to 2. Analytical grade ethyl acetate (Sigma Aldrich) was used to perform liquid-liquid extraction. The upper phases were collected, and the residual moisture was removed using magnesium-carbonate (Sigma Aldrich) after combining the extracts. Finally, the solvent was removed from the ethyl acetate soluble extract (Fraction A252E) using a Büchi Rotavapor R-100 bench-top evaporator system.

### Preparative scale High Performance Liquid Chromatography (HPLC) -reverse phase

Waters AutoPurification system was used for the preparative HPLC, which consisted of the 2545 Binary Gradient Module, 2767 Injector/Collector Sample Manager, System Fluidics Organizer, and 2998 Photodiode Array Detector. The samples were dissolved in DMSO (Sigma Aldrich) at a very high concentration. They were injected onto a Phenomenex Luna Phenyl-Hexyl 100A (5mL), (250×50 mm, 10µm) or Phenomenex Luna C18(2) 100A, (250×21.5 mm, 5 µm) columns to separate components. The flow rate was 42.48 ml/min, and the eluents were: “Eluent A” was analytical grade Water (Sigma Aldrich) with 0.05% Formic acid (Sigma Aldrich), and “Eluent B” was analytical grade Acetonitrile (Sigma Aldrich) with 0.05% Formic acid. The gradient was the following: 0 min 95% A, 1 min 75% A, 86.76 min 70% A, 91.67 min 5% A, 96.67 min 5% A, and 100 min 95% A. The fractions were collected continuously between 5 and 25 minutes. In case the PDA detector triggered a fraction at a specific absorbance, the components were collected. The system was controlled by the Waters MassLynx 4.1 software with the FractionLynx add-on.

### Liquid-liquid extraction of the lipophilic part

50g of Fraction A250 was resuspended in water. The pH was adjusted to 2 with Hydrochloric acid (Sigma Aldrich) and was extracted twice with ethyl acetate (Sigma Aldrich). The extract was dried with magnesium carbonate (Sigma-Aldrich) and filtered with Buchi glass filter utilizing a quantitative grade filter paper (Millipore). The solvent was removed under vacuum at 50^ο^C using a Buchi Rotavapor R-100 bench-top evaporator system.

### Flash chromatography

A Teledyne Combiflash Companion Chromatography System with RedySep Rf chromatography column prepacked with 80 grams of normal phase silica was used to separate the components of the concentrated fractions. Eluent A was analytical grade Hexene (Sigma Aldrich), and eluent B was analytical grade Ethyl Acetate. The flow rate was set to 60 ml/min, the gradient was 2min 100% A, 27min 100% B for 4 minutes, then 100% A for 3 min. The sample was dissolved in methanol at very high concentration. The fractions were collected continuously from the beginning of the run. Selected fractions were dried with a Genevac EZ-2 Plus Personal in a 20 ml sample vial.

### Preparative scale HPLC purification

Waters AutoPurification system consisted of the 2545 Binary Gradient Module, 2767 Injector/Collector Sample Manager, System Fluidics Organizer, and 2998 Photodiode Array Detector was used to isolate fractions of A252. Samples were dissolved in dimethoxy sulfoxide (Sigma Aldrich) at very high concentration and injected up to 5 mL volume on to a Phenomenex Luna Phenyl-Hexyl 100A, (250×50 mm, 10u) or Phenomenex Luna C18(2) 100A, (250×21.5 mm, 5 µm) columns to separate components. The flow rate was 42.48 ml/min. The eluents were the following: “Eluent A” was analytical grade Water (Sigma Aldrich) with 0.05% Formic acid (Sigma Aldrich), and “Eluent B” was analytical grade Acetonitrile (Sigma Aldrich) with 0.05% Formic. The gradient was the following: 0 min 95% A, 1 min 75% A, 86.76 min 70% A, 91.67 min 5% A, 96.67 min 5% A and 100 min 95% A. The factions were collected either continuously between 5 and 25 minutes of the run, or the collection was triggered by the PDA detector at a specified absorbance in case a specific component was collected. The system was controlled by the Waters MassLynx 4.1 software with FractionLynx add-on.

### Mass spectrometry (LC-MS/MS)

A Thermo Scientific TSQ Vantage Triple Quadrupole mass spectrometer to quantify the isolated inhibitor in the fermentation broth. The sample was eluted from the Thermo Accela HPLC system equipped with a Thermo Scientific Aquasil C18 Column (100 x 2.1 mm, 3 μm). The flow rate was 100 µl/min and the eluents were analytical grade Water (Sigma Aldrich) with 0.05% Formic acid (Sigma Aldrich) as eluent A, and analytical grade Acetonitrile (Sigma Aldrich) with 0.05% Formic acid as Eluent B. I used the synthetic compound with verified purity for tuning and calibration. The main parameters of the detector were Capillary temperature 320°C, vaporizer temperature 300°C, Sheath gas pressure 30 psi, Auxiliary gas flow 25, Spray voltage 3000 V. The parent ion of the inhibitor was 347.966Da, and the most abundant daughter ions were 166.975Da, 181.971Da, 207.949Da, and 225.969Da.

### Peptide-based kinase assay

The radiometric peptide-based kinase assay was performed to validate the results and determine the IC50 of the inhibitors using the protocol described in^48^. The recombinant His-tagged (N-terminus) DYRK1A (127-485aa) and PIM1 kinases (313aa) were purified from bacterial BL21 cells using Ni-NTA beads and assayed against the peptide sequences KKISGRLSPIMTEQ and RSRHSSYPAGT (gift from University of Dundee) for DYRK1A and PIM1 respectively at a final concentration of 10mM in the assay buffer containing 25mM HEPES, pH 6.8, 5mM MgCl_2_, 0.5mM DTT and 1mg/ml BSA. The [g-^32^P] ATP for the assay was purchased from Perkin Elmer (#BLU502A250UC). The specific activity of [g-^32^P] ATP was determined by dissolving and spiking non-radioactive ‘cold’ ATP (Sigma) in assay buffer using [g-^32^P] ATP to produce radioactivity of 1 x 10^5^ to 1 x 10^6^ cpm per nmol. The kinase assay done either with or without inhibitors was performed at 30°C with a 10 mins incubation period. The reaction mixture (15ml) was then spotted onto 2×2 P81 phosphocellulose papers (Whatman) that bind the peptide substrate. The excess ATP was then washed from the P81 papers using 75mM phosphoric acid using a wire mesh basket three times followed by a final wash using acetone. The P81 papers were dried and measured for radioactivity with a scintillation counter (GMI). The data were then plotted using GraphPad Prism.

### Chemical synthesis of CSH-4044 (2,5-Dihydroxy-4-methoxy-phenyl)-(6-hydroxy-5-methoxy-benzothiazol-2-yl)-methanone

#### 4-Hydroxy-3-methoxy-phenyl)-thiourea (1)

A mixture of 1.39 g (10.0 mmol) 4-amino-2-methoxyphenol, 2.18 ml of water, and 0.78 ml hydrochloric acid (37%) was stirred at 80°C in a closed vessel for 1 hour, then 0,76 g (10 mmol) of ammonium thiocyanate was added, and the stirring was continued at 120°C for 1 day. The reaction mixture was diluted with water, the precipitate was filtered out, washed with water and a small amount of acetone, then dried in air. Thus, 1.48 g of the title compound was obtained (yield 75%). LCMS Anal. calculated for C8H10N2O2S: 198.25, found m/z: 197 [M-H]-, 199 [M+H]+, tR = 0.46 and 1.21 min; peak area 99%.

#### 2-Amino-6-hydroxy-5-methoxy-benzothiazole (2)

To the solution of 1.43g (7.21 mmol) (4-hydroxy-3-methoxy-phenyl)-thiourea (1) in 42ml acetic acid, 2.84g (7.28mmol) benzyltrimethylammonium tribromide was added in portions, then the mixture was stirred at room temperature for 1 hour. The precipitated hydrogenbromide salt was filtered out, washed with acetic acid, and diisopropyl ether. This solid was dissolved in a saturated aqueous solution of sodium bicarbonate, then extracted three times with ethyl acetate. The organic layers were combined, washed with brine, dried over anhydrous sodium sulfate, filtered, and the solvent was evaporated. The residue solidified under diisopropyl ether. Thus, 1.05 g 2-amino-5-methoxy-benzothiazol-6-ol (2) was obtained (yield: 74%). Melting point: 183-185°C. 1H NMR (300 MHz, DMSO-d6): δ 8.59 (s, 1H), 7.05 (s, 2H), 7.01 (s, 1H), 6.93 (s, 1H), 3.76 (s, 3H). LCMS Anal. calculated for C8H8N2O2S: 196.23, found m/z: 197 [M+H]+, tR = 0.46 and 1.42 min; peak area 97%.

#### Bis(4-hydroxi-5-methoxyanilin-2-yl)disulfide (3)

A mixture of 3000 mg of potassium hydroxide, 9ml of water, and 1.00g (5.09 mmol) 2-amino-6-hydroxy-5-methoxy-benzothiazole was refluxed for 6 hours. The cooled solution was neutralized with acetic acid, and then sodium hydrogen carbonate solution was added. The solution was extracted three times with ethyl acetate. The organic phases were combined, washed with brine, dried over anhydrous sodium sulfate, filtered, and the solvent was evaporated in a vacuum. The residue solidified under diisopropyl ether. Thus, 632 mg of the title compound was obtained (yield 81%). 1H NMR (300 MHz, DMSO-d6): δ 8.15 (s, 2H), 6.55 (s, 2H), 6.35 (s, 2H), 4.83 (s, 4H), 3.70 (s, 6H). LCMS Anal. calculated for C14H16N2O4S2: 340.42, found m/z: 339 [M+H]-, 341 [M+H]+, tR = 0.46, and 2.07 min; peak area 96%.

#### (2,5-Dihydroxy-4-methoxy-phenyl)-oxo-acetic acid ethyl ester (4)

1.4g (10 mmol) of 2,5-dihydroxyanisole and 1.38g (10.1 mmol) of ethyl oxalyl chloride were dissolved in 100 ml of anhydrous dichloromethane, stirred at −15°C. Then 12 ml of 1M solution of titanium tetrachloride in dichloromethane was added dropwise to the reaction mixture at −15 °C for 30 minutes, and the stirring was continued for a further 2 hours at −15 °C. 150 ml 1M hydrochloric acid was added to the reaction mixture, then stirred at room temperature for 1 hour. The two layers were separated, the aqueous layer was extracted two times with dichloromethane, the organic layers were combined, washed with 100ml water, then with 50ml saturated sodium hydrogen carbonate solution and 100ml water, the organic layer was separated and dried over sodium sulfate, filtered and evaporated in vacuum. The residue solidified under hexane. Thus 1.072g of title compound was obtained (yield 39%). Melting point: 131-132°C. 1H NMR (300 MHz, DMSO-d6): δ 10.60 (s, 1H), 9.00 (s, 1H), 7.04 (s, 1H), 6.46 (s, 1H), 4.31-4.24 (q, J = 7.2 Hz, 2H), 1.30-1.25 (t, J = 7.2 Hz, 3H). LCMS Anal. calculated for C11H12O6: 240.21, found m/z: 239 [M+H]-, 241 [M+H]+, tR = 2.99 min; peak area 99%.

#### (2,5-Dihydroxy-4-methoxy-phenyl)-(6-hydroxy-5-methoxy-benzothiazol-2-yl)-methanone (5)

408 mg (1.2 mmol) of disulfide (3) and 312 mg (1.2 mmol) of triphenylphosphine were stirred in aqueous ethyl alcohol (2.4 ml of water and 2.4 ml of ethyl alcohol) under argon atmosphere for 1 hour at room temperature. Then 652 mg (2.4 mmol) of ester (4) was added, argon was passed through the mixture for 10 minutes, the vessel was closed, and it was put into a microwave oven for 2.5 hours at 130 °C. The precipitate was filtered at room temperature, washed with ethyl alcohol and diisopropyl ether, then dried in air. The crude product was purified by preparative HPLC. Thus, 350 mg of the title compound was obtained (yield 42%). 1H NMR (300 MHz, DMSO-d6): δ 12.46 (s, 1H), 10.12 (b, 1H), 8.96 (s, 1H), 8.66 (s, 1H), 7.72 (s, 1H), 7.51 (s, 1H), 6.60 (s, 1H), 3.93 (s, 3H), 3.90 (s, 3H). LCMS Anal. calculated for C16H13NO6S: 347.35, found m/z: 346 [M+H], 348 [M+H]+, tR = 3.54 min; peak area 99%.

### Protein expression, purification and crystallization

Human PIM1 (PIM1) (1-313aa) was cloned into the pET28 expression vector with a N-terminal 6XHis and SUMO tags, followed by a TEV protease cleavage site. The protein was expressed in *in E.coli (BL21 DE3)* for 16 hours at 20° C. Cells were resuspended and lysed by sonication in 50 mM Tris pH=8.0, 100 mM NaCl, 40 mM Imidazole and 2 mM CaCl_2_. Protein was initially purified using Ni-NTA agarose resin (Qiagene) followed by on column TEV protease cleavage. Further purification was carried out using a Q-HP column (Cytiva) followed by a final purification using a HiLoad Superdex 75 column (Cytiva). Protein was stored in the following buffer: 20 mM Tris pH=8.0, 100 mM NaCl, 2 mM MgCl_2_, 2 mM CaCl_2_ 0.5 mM TCEP, and 5 mM DTT at 5 mg/ml. The protein was crystallized using the vapor diffusion sitting drop method by mixing 500nL of the protein with 500nL of 0.2 M NaCl, 0.2 M CaCl_2_, 0.1 M tri-Sodium-citrate pH=5.5 and 1 M di-ammonium hydrogen phosphate precipitant solution at 18° C. Crystals were then soaked in 0.2 M NaCl, 0.2 M CaCl_2_, 0.1 M tri-Sodium-citrate pH=5.5, and 1 M di-ammonium hydrogen phosphate supplemented with 10 mM CSH-4044 overnight. Crystals were cryoprotected by transferring them briefly to a solution containing 0.2 M NaCl, 0.2 M CaCl_2_, 0.1 M tri-sodium-citrate pH=5.5, 1 M di-ammonium hydrogen phosphate, and 30% glycerol before flash freezing in liquid nitrogen.

### Data collection and structure determination

Data were collected to 1.95 Å resolution at beamline 8.2.1 at the Berkeley Center for Structural Biology (BCSB) at the Advanced Light Source (ALS). Diffraction data were indexed, integrated, and scaled using autoPROC^49^. The structure was solved by molecular replacement in PHASER^50^ using the Apo PIM1 structure as a search model (PDB: 1XQZ)^25^. The molecular replacement solution was rigid-body refined in PHENIX followed by simulated annealing refinement prior to manual correction in COOT^51^. The final refinement of the model was done using PHENIX^52^ and validated by Molprobity^53^. X-ray data collection and refinement statistics for PIM1-CSH-4044 structure are summarized in Table-2. The atomic coordinates and structure factors for PIM1-CSH-4044 structure have been deposited in the Protein Data Bank under accession code 9YKI.

### Structure determination of CSH-4044 using X-ray crystallography

The structure of the isolated compound, CSH-4044 was determined at New York University, New York (NYU) by Dr. Chunhuao Tony Hu. A red needle-like crystal with the size of 0.01 x 0.05 x 0.28 mm^3^ was selected for geometry and intensity data collection with a Bruker SMART APEXII CCD area detector on a D8 goniometer at 100 K. The temperature during the data collection was controlled with an Oxford Cryosystems Series 700 plus instrument. Preliminary lattice parameters and orientation matrices were obtained from three sets of frames. Data were collected using graphite-monochromated and 0.5 mm-MonoCap-collimated Mo-Ka radiation (λ = 0.71073 Å) with the ω scan method. Data was processed with the INTEGRATE program of the APEX2 software for reduction and cell refinement. Multi-scan absorption corrections were applied by using the SCALE program for the area detector. The structure was solved by the direct method and refined on F2 (SHELXTL). Non-hydrogen atoms were refined with anisotropic displacement parameters, and hydrogen atoms on carbons and oxygens were placed in idealized positions (C-H = 0.95-0.98 Å and O-H = 0.84 Å) and included as riding with Uiso(H) = 1.2 or 1.5 Ueq(non-H).

### Cell-based assays

The cell-lines MiaPaca-2 and SH-SY5Y was obtained from ATCC. After treatment with CSH-4044 or DMSO, whole cell lysates were harvested and resuspended in RIPA buffer [25 mM Tris, pH 7.4, 150 mM NaCl, 1% Triton X 100, 0.5% sodium deoxycholate, 0.1% sodium dodecyl sulfate, protease inhibitor cocktail (Sigma, Cat. No. 4693159001), and phosphatase inhibitor cocktail (Sigma, Cat. No. 4906845001)]. Quantification of protein concentration was done using the Pierce™ BCA Protein Assay kit (ThermoFisher, Catalog #23225). Equal amounts of lysate were denatured and loaded onto a 10% SDS-PAGE gel. Antibody blocking was done with 5% milk in TBST (19mM Tris base, NaCl 137 mM, KCl 2.7mM and 0.1% Tween-20) for 1 hour at room temperature. Blots were incubated with the primary antibody pBAD Ser112 (Cell Signaling Technologies, Cat #9291), pTau T212, pTau S202/T205, pTau T231 and Total Tau (Thermofisher Scientific, Cat #44-740G, #MN1020, #MA5-12808, #MN1040) overnight at 4°C. Membranes were washed three times at room temperature (10 mins each) before they were incubated with secondary antibodies for one hour at room temperature. HRP goat anti-mouse (Bio-Rad; Cat. No. 1706516) at 1:50,000 was used for tubulin blots while HRP goat anti-rabbit (Abcam, Cat. No. ab6721) at 1:30,000 was used for all other primary antibodies. Membranes were washed 3 times again (15 min each) and developed using SuperSignal™ West Dura Extended Duration Substrate (ThermoFisher, Catalog #34075) and BioExcell autoradiographic film (Worldwide Medical, Catalog # 41101001).

## Acknowledgements

We thank Dr. Chunhua Hu and the NYU Molecular Design Institute for assistance with data collection and structure determination of CS4044. We thank Dr. James D. Watson for his encouragement and his generous personal donation that enabled the purchase of the preparative-scale HPLC system used for fractionating the extract and isolating the inhibitor compound. His support was instrumental to the success of this project.

## REFERENCES

(1) Zhurakivska, K.; Troiano, G.; Caponio, V. C. A.; Dioguardi, M.; Arena, C.; Lo Muzio, L. The Effects of Adjuvant Fermented Wheat Germ Extract on Cancer Cell Lines: A Systematic Review. Nutrients 2018, 10 (10). 10.3390/nu10101546.

(2) Imir, N. G.; Aydemir, E.; Şimşek, E. Mechanism of the Anti-Angiogenic Effect of Avemar on Tumor Cells. Oncol. Lett. 2018, 15 (2), 2673–2678. 10.3892/ol.2017.7604.

(3) Mueller, T.; Jordan, K.; Voigt, W. Promising Cytotoxic Activity Profile of Fermented Wheat Germ Extract (Avemar®) in Human Cancer Cell Lines. J. Exp. Clin. Cancer Res. CR 2011, 30, 42. 10.1186/1756-9966-30-42.

(4) Judson, P. L.; Al Sawah, E.; Marchion, D. C.; Xiong, Y.; Bicaku, E.; Bou Zgheib, N.; Chon, H. S.; Stickles, X. B.; Hakam, A.; Wenham, R. M.; Apte, S. M.; Gonzalez-Bosquet, J.; Chen, D.-T.; Lancaster, J. M. Characterizing the Efficacy of Fermented Wheat Germ Extract against Ovarian Cancer and Defining the Genomic Basis of Its Activity. Int. J. Gynecol. Cancer Off. J. Int. Gynecol. Cancer Soc. 2012, 22 (6), 960– 967. 10.1097/IGC.0b013e318258509d.

(5) Yang, M.-D.; Chang, W.-S.; Tsai, C.-W.; Wang, M.-F.; Chan, Y.-C.; Chan, K.-C.; Lu, M.-C.; Kao, A.-W.; Hsu, C.-M.; Bau, D.-T. Inhibitory Effects of AVEMAR on Proliferation and Metastasis of Oral Cancer Cells. Nutr. Cancer 2016, 68 (3), 473– 480. 10.1080/01635581.2016.1153668.

(6) Telekes, A.; Hegedus, M.; Chae, C.-H.; Vékey, K. Avemar (Wheat Germ Extract) in Cancer Prevention and Treatment. Nutr. Cancer 2009, 61 (6), 891–899. 10.1080/01635580903285114.

(7) Hidvégi, M.; Ráso, E.; Tömösközi-Farkas, R.; Paku, S.; Lapis, K.; Szende, B. Effect of Avemar and Avemar + Vitamin C on Tumor Growth and Metastasis in Experimental Animals. Anticancer Res. 1998, 18 (4A), 2353–2358.

(8) Nawijn, M. C.; Alendar, A.; Berns, A. For Better or for Worse: The Role of Pim Oncogenes in Tumorigenesis. Nat. Rev. Cancer 2011, 11 (1), 23–34. 10.1038/nrc2986.

(9) Fernández-Martínez, P.; Zahonero, C.; Sánchez-Gómez, P. DYRK1A: The Double-Edged Kinase as a Protagonist in Cell Growth and Tumorigenesis. Mol. Cell. Oncol. 2015, 2 (1), e970048. 10.4161/23723548.2014.970048.

(10) Bencze, G.; Bencze, S.; Rivera, K. D.; Watson, J. D.; Hidvegi, M.; Orfi, L.; Tonks, N. K.; Pappin, D. J. Mito-Oncology Agent: Fermented Extract Suppresses the Warburg Effect, Restores Oxidative Mitochondrial Activity, and Inhibits in Vivo Tumor Growth. Sci. Rep. 2020, 10 (1), 14174. 10.1038/s41598-020-71118-3.

(11) Kumar, A.; Mandiyan, V.; Suzuki, Y.; Zhang, C.; Rice, J.; Tsai, J.; Artis, D. R.; Ibrahim, P.; Bremer, R. Crystal Structures of Proto-Oncogene Kinase Pim1: A Target of Aberrant Somatic Hypermutations in Diffuse Large Cell Lymphoma. J. Mol. Biol. 2005, 348 (1), 183–193. 10.1016/j.jmb.2005.02.039.

(12) Jacobs, M. D.; Black, J.; Futer, O.; Swenson, L.; Hare, B.; Fleming, M.; Saxena, K. Pim-1 Ligand-Bound Structures Reveal the Mechanism of Serine/Threonine Kinase Inhibition by LY294002. J. Biol. Chem. 2005, 280 (14), 13728–13734. 10.1074/jbc.M413155200.

(13) Aho, T. L. T.; Sandholm, J.; Peltola, K. J.; Mankonen, H. P.; Lilly, M.; Koskinen, P. J. Pim-1 Kinase Promotes Inactivation of the pro-Apoptotic Bad Protein by Phosphorylating It on the Ser ^112^ Gatekeeper Site. FEBS Lett. 2004, 571 (1–3), 43– 49. 10.1016/j.febslet.2004.06.050.

(14) Yan, B.; Zemskova, M.; Holder, S.; Chin, V.; Kraft, A.; Koskinen, P. J.; Lilly, M. The PIM-2 Kinase Phosphorylates BAD on Serine 112 and Reverses BAD-Induced Cell Death. J. Biol. Chem. 2003, 278 (46), 45358–45367. 10.1074/jbc.M307933200.

(15) Li, Y.-Y.; Popivanova, B. K.; Nagai, Y.; Ishikura, H.; Fujii, C.; Mukaida, N. Pim-3, a Proto-Oncogene with Serine/Threonine Kinase Activity, Is Aberrantly Expressed in Human Pancreatic Cancer and Phosphorylates Bad to Block Bad-Mediated Apoptosis in Human Pancreatic Cancer Cell Lines. Cancer Res. 2006, 66 (13), 6741–6747. 10.1158/0008-5472.can-05-4272.

(16) Popivanova, B. K.; Li, Y.-Y.; Zheng, H.; Omura, K.; Fujii, C.; Tsuneyama, K.; Mukaida, N. Proto-Oncogene, Pim-3 with Serine/Threonine Kinase Activity, Is Aberrantly Expressed in Human Colon Cancer Cells and Can Prevent Bad-Mediated Apoptosis. Cancer Sci. 2007, 98 (3), 321–328. 10.1111/j.1349-7006.2007.00390.x.

(17) Liu, B.; Wang, Z.; Li, H.-Y.; Zhang, B.; Ping, B.; Li, Y.-Y. Pim-3 Promotes Human Pancreatic Cancer Growth by Regulating Tumor Vasculogenesis. Oncol. Rep. 2014, 31 (6), 2625–2634. 10.3892/or.2014.3158.

(18) Ryoo, S.-R.; Jeong, H. K.; Radnaabazar, C.; Yoo, J.-J.; Cho, H.-J.; Lee, H.-W.; Kim, I.-S.; Cheon, Y.-H.; Ahn, Y. S.; Chung, S.-H.; Song, W.-J. DYRK1A-Mediated Hyperphosphorylation of Tau. A Functional Link between Down Syndrome and Alzheimer Disease. J. Biol. Chem. 2007, 282 (48), 34850–34857. 10.1074/jbc.M707358200.

(19) Yin, X.; Jin, N.; Shi, J.; Zhang, Y.; Wu, Y.; Gong, C.-X.; Iqbal, K.; Liu, F. Dyrk1A Overexpression Leads to Increase of 3R-Tau Expression and Cognitive Deficits in Ts65Dn Down Syndrome Mice. Sci. Rep. 2017, 7 (1), 619. 10.1038/s41598-017-00682-y.

(20) Chaves, J. C. S.; Machado, F. T.; Almeida, M. F.; Bacovsky, T. B.; Ferrari, M. F. R. microRNAs Expression Correlates with Levels of APP, DYRK1A, Hyperphosphorylated Tau and BDNF in the Hippocampus of a Mouse Model for Down Syndrome during Ageing. Neurosci. Lett. 2020, 714, 134541. 10.1016/j.neulet.2019.134541.

(21) Fajka-Boja, R.; Hidvégi, M.; Shoenfeld, Y.; Ion, G.; Demydenko, D.; Tömösközi-Farkas, R.; Vizler, C.; Telekes, A.; Resetar, A.; Monostori, E. Fermented Wheat Germ Extract Induces Apoptosis and Downregulation of Major Histocompatibility Complex Class I Proteins in Tumor T and B Cell Lines. Int. J. Oncol. 2002, 20 (3), 563–570.

(22) Marcsek, Z.; Kocsis, Z.; Jakab, M.; Szende, B.; Tompa, A. The Efficacy of Tamoxifen in Estrogen Receptor-Positive Breast Cancer Cells Is Enhanced by a Medical Nutriment. Cancer Biother. Radiopharm. 2004, 19 (6), 746–753. 10.1089/cbr.2004.19.746.

(23) Comin-Anduix, B.; Boros, L. G.; Marin, S.; Boren, J.; Callol-Massot, C.; Centelles, J. J.; Torres, J. L.; Agell, N.; Bassilian, S.; Cascante, M. Fermented Wheat Germ Extract Inhibits Glycolysis/Pentose Cycle Enzymes and Induces Apoptosis through Poly(ADP-Ribose) Polymerase Activation in Jurkat T-Cell Leukemia Tumor Cells. J. Biol. Chem. 2002, 277 (48), 46408–46414. 10.1074/jbc.M206150200.

(24) Bullock, A. N.; Debreczeni, J.; Amos, A. L.; Knapp, S.; Turk, B. E. Structure and Substrate Specificity of the Pim-1 Kinase. J. Biol. Chem. 2005, 280 (50), 41675– 41682. 10.1074/jbc.M510711200.

(25) Qian, K. C.; Wang, L.; Hickey, E. R.; Studts, J.; Barringer, K.; Peng, C.; Kronkaitis, A.; Li, J.; White, A.; Mische, S.; Farmer, B. Structural Basis of Constitutive Activity and a Unique Nucleotide Binding Mode of Human Pim-1 Kinase. J. Biol. Chem. 2005, 280 (7), 6130–6137. 10.1074/jbc.M409123200.

(26) Bullock, A. N.; Russo, S.; Amos, A.; Pagano, N.; Bregman, H.; Debreczeni, J. É.; Lee, W. H.; Delft, F. von; Meggers, E.; Knapp, S. Crystal Structure of the PIM2 Kinase in Complex with an Organoruthenium Inhibitor. PLoS ONE 2009, 4 (10), e7112. 10.1371/journal.pone.0007112.

(27) Qian, K. C.; Studts, J.; Wang, L.; Barringer, K.; Kronkaitis, A.; Peng, C.; Baptiste, A.; LaFrance, R.; Mische, S.; Farmer, B. Expression, Purification, Crystallization and Preliminary Crystallographic Analysis of Human Pim-1 Kinase. Acta Crystallograph. Sect. F Struct. Biol. Cryst. Commun. 2005, 61 (Pt 1), 96–99. 10.1107/S1744309104029963.

(28) Razmazma, H.; Ebrahimi, A.; Hashemi, M. Structural Insights for Rational Design of New PIM-1 Kinase Inhibitors Based on 3,5-Disubstituted Indole Derivatives: An Integrative Computational Approach. Comput. Biol. Med. 2020, 118, 103641. 10.1016/j.compbiomed.2020.103641.

(29) Bogusz, J.; Zrubek, K.; Rembacz, K. P.; Grudnik, P.; Golik, P.; Romanowska, M.; Wladyka, B.; Dubin, G. Structural Analysis of PIM1 Kinase Complexes with ATP-Competitive Inhibitors. Sci. Rep. 2017, 7 (1), 13399. 10.1038/s41598-017-13557-z.

(30) Sever, R.; Brugge, J. S. Signal Transduction in Cancer. Cold Spring Harb. Perspect. Med. 2015, 5 (4). 10.1101/cshperspect.a006098.

(31) Bhullar, K. S.; Lagarón, N. O.; McGowan, E. M.; Parmar, I.; Jha, A.; Hubbard, B. P.; Rupasinghe, H. P. V. Kinase-Targeted Cancer Therapies: Progress, Challenges and Future Directions. Mol. Cancer 2018, 17 (1), 48. 10.1186/s12943-018-0804-2.

(32) Bachmann, M.; Hennemann, H.; Xing, P. X.; Hoffmann, I.; Möröy, T. The Oncogenic Serine/Threonine Kinase Pim-1 Phosphorylates and Inhibits the Activity of Cdc25C-Associated Kinase 1 (C-TAK1): A NOVEL ROLE FOR Pim-1 AT THE G _2_ /M CELL CYCLE CHECKPOINT. J. Biol. Chem. 2004, 279 (46), 48319–48328. 10.1074/jbc.M404440200.

(33) Winn, L. M.; Lei, W.; Ness, S. A. Pim-1 Phosphorylates the DNA Binding Domain of c-Myb. Cell Cycle Georget. Tex 2003, 2 (3), 258–262.

(34) Rainio, E.-M.; Sandholm, J.; Koskinen, P. J. Cutting Edge: Transcriptional Activity of NFATc1 Is Enhanced by the Pim-1 Kinase. J. Immunol. Baltim. Md 1950 2002, 168 (4), 1524–1527. 10.4049/jimmunol.168.4.1524.

(35) Wang, J.; Kim, J.; Roh, M.; Franco, O. E.; Hayward, S. W.; Wills, M. L.; Abdulkadir, S. A. Pim1 Kinase Synergizes with C-MYC to Induce Advanced Prostate Carcinoma. Oncogene 2010, 29 (17), 2477–2487. 10.1038/onc.2010.10.

(36) Fox, C. J.; Hammerman, P. S.; Cinalli, R. M.; Master, S. R.; Chodosh, L. A.; Thompson, C. B. The Serine/Threonine Kinase Pim-2 Is a Transcriptionally Regulated Apoptotic Inhibitor. Genes Dev. 2003, 17 (15), 1841–1854. 10.1101/gad.1105003.

(37) Koike, N.; Maita, H.; Taira, T.; Ariga, H.; Iguchi-Ariga, S. M. M. Identification of Heterochromatin Protein 1 (HP1) as a Phosphorylation Target by Pim-1 Kinase and the Effect of Phosphorylation on the Transcriptional Repression Function of HP1 ^1^. FEBS Lett. 2000, 467 (1), 17–21. 10.1016/S0014-5793(00)01105-4.

(38) Leverson, J. D.; Koskinen, P. J.; Orrico, F. C.; Rainio, E. M.; Jalkanen, K. J.; Dash, A. B.; Eisenman, R. N.; Ness, S. A. Pim-1 Kinase and P100 Cooperate to Enhance c-Myb Activity. Mol. Cell 1998, 2 (4), 417–425. 10.1016/s1097-2765(00)80141-0.

(39) Mochizuki, T.; Kitanaka, C.; Noguchi, K.; Muramatsu, T.; Asai, A.; Kuchino, Y. Physical and Functional Interactions between Pim-1 Kinase and Cdc25A Phosphatase: IMPLICATIONS FOR THE Pim-1-MEDIATED ACTIVATION OF THE c-Myc SIGNALING PATHWAY. J. Biol. Chem. 1999, 274 (26), 18659–18666. 10.1074/jbc.274.26.18659.

(40) Wang, Z.; Bhattacharya, N.; Mixter, P. F.; Wei, W.; Sedivy, J.; Magnuson, N. S. Phosphorylation of the Cell Cycle Inhibitor p21Cip1/WAF1 by Pim-1 Kinase. Biochim. Biophys. Acta 2002, 1593 (1), 45–55. 10.1016/s0167-4889(02)00347-6.

(41) Mukaida, N.; Wang, Y.-Y.; Li, Y.-Y. Roles of Pim-3, a Novel Survival Kinase, in Tumorigenesis. Cancer Sci. 2011, 102 (8), 1437–1442. 10.1111/j.1349-7006.2011.01966.x.

(42) Xu, D.; Cobb, M. G.; Gavilano, L.; Witherspoon, S. M.; Williams, D.; White, C. D.; Taverna, P.; Bednarski, B. K.; Kim, H. J.; Baldwin, A. S.; Baines, A. T. Inhibition of Oncogenic Pim-3 Kinase Modulates Transformed Growth and Chemosensitizes Pancreatic Cancer Cells to Gemcitabine. Cancer Biol. Ther. 2013, 14 (6), 492–501. 10.4161/cbt.24343.

(43) Abbassi, R.; Johns, T. G.; Kassiou, M.; Munoz, L. DYRK1A in Neurodegeneration and Cancer: Molecular Basis and Clinical Implications. Pharmacol. Ther. 2015, 151, 87–98. 10.1016/j.pharmthera.2015.03.004.

(44) Becker, W.; Soppa, U.; Tejedor, F. J. DYRK1A: A Potential Drug Target for Multiple Down Syndrome Neuropathologies. CNS Neurol. Disord. Drug Targets 2014, 13 (1), 26–33. 10.2174/18715273113126660186.

(45) Demuro, S.; Di Martino, R. M. C.; Ortega, J. A.; Cavalli, A. GSK-3β, FYN, and DYRK1A: Master Regulators in Neurodegenerative Pathways. Int. J. Mol. Sci. 2021, 22 (16), 9098. 10.3390/ijms22169098.

(46) Aranda, S.; Laguna, A.; de la Luna, S. DYRK Family of Protein Kinases: Evolutionary Relationships, Biochemical Properties, and Functional Roles. FASEB J. Off. Publ. Fed. Am. Soc. Exp. Biol. 2011, 25 (2), 449–462. 10.1096/fj.10-165837.

(47) Asati, V.; Mahapatra, D. K.; Bharti, S. K. PIM Kinase Inhibitors: Structural and Pharmacological Perspectives. Eur. J. Med. Chem. 2019, 172, 95–108. 10.1016/j.ejmech.2019.03.050.

(48) Hastie, C. J.; McLauchlan, H. J.; Cohen, P. Assay of Protein Kinases Using Radiolabeled ATP: A Protocol. Nat. Protoc. 2006, 1 (2), 968–971. 10.1038/nprot.2006.149.

(49) Vonrhein, C.; Flensburg, C.; Keller, P.; Sharff, A.; Smart, O.; Paciorek, W.; Womack, T.; Bricogne, G. Data Processing and Analysis with the autoPROC Toolbox. Acta Crystallogr. D Biol. Crystallogr. 2011, 67 (Pt 4), 293–302. 10.1107/S0907444911007773.

(50) McCoy, A. J.; Grosse-Kunstleve, R. W.; Adams, P. D.; Winn, M. D.; Storoni, L. C.; Read, R. J. *Phaser* Crystallographic Software. J. Appl. Crystallogr. 2007, 40 (4), 658–674. 10.1107/S0021889807021206.

(51) Emsley, P.; Lohkamp, B.; Scott, W. G.; Cowtan, K. Features and Development of *Coot*. Acta Crystallogr. D Biol. Crystallogr. 2010, 66 (4), 486–501. 10.1107/S0907444910007493.

(52) Adams, P. D.; Afonine, P. V.; Bunkóczi, G.; Chen, V. B.; Davis, I. W.; Echols, N.; Headd, J. J.; Hung, L.-W.; Kapral, G. J.; Grosse-Kunstleve, R. W.; McCoy, A. J.; Moriarty, N. W.; Oeffner, R.; Read, R. J.; Richardson, D. C.; Richardson, J. S.; Terwilliger, T. C.; Zwart, P. H. *PHENIX* : A Comprehensive Python-Based System for Macromolecular Structure Solution. Acta Crystallogr. D Biol. Crystallogr. 2010, 66 (2), 213–221. 10.1107/S0907444909052925.

(53) Chen, V. B.; Arendall, W. B.; Headd, J. J.; Keedy, D. A.; Immormino, R. M.; Kapral, G. J.; Murray, L. W.; Richardson, J. S.; Richardson, D. C. *MolProbity* : All-Atom Structure Validation for Macromolecular Crystallography. Acta Crystallogr. D Biol. Crystallogr. 2010, 66 (1), 12–21. 10.1107/S0907444909042073.

