## Supplemental figures for "Identification and Validation of an inhibitor of the protein kinases PIM and DYRK"

### Supplementary Fig 1

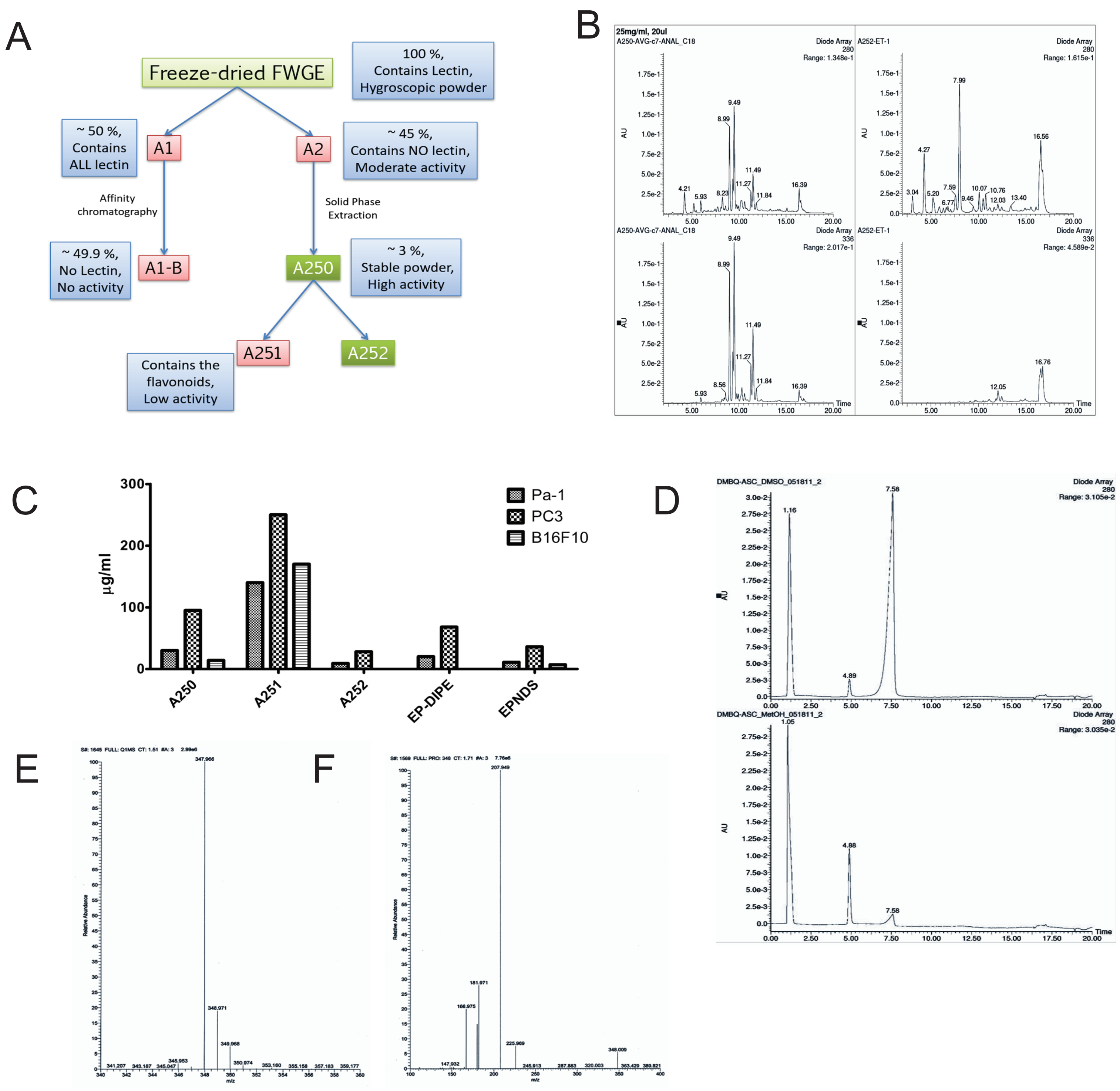


**Figure S1. Enrichment of Fraction A250 from Fermented wheat germ extract (FWGE)**

**(A)** Isolation diagram of Fraction A252 from freeze-dried FWGE. **(B)** HPLC-UV chromatogram of Fraction A250 (left) and A252-E (right) recorded at 280 and 336 nm, the latter corresponding to the characteristic absorbance of flavonoids. **(C)** In vitro activity of Fraction A250 and its subfractions on Pa-1, B16F10 and PC3 cells. **(D)** Standard 2,6-DMBQ (at 7.58 min) with ascorbic acid in DMSO (upper panel) and methanol (lower panel). In methanol, 2,6-DMBQ was reduced to the hydroquinone form, whereas no reduction occurred in DMSO. **(E)** Electrospray ionization (ESI) mass spectrum of the inhibitor, showing the molecular ion and isotopic distribution. **(F)** ESI-MS/MS spectrum of the inhibitor, highlighting the major fragment (daughter) ions.

### Supplementary Fig 2

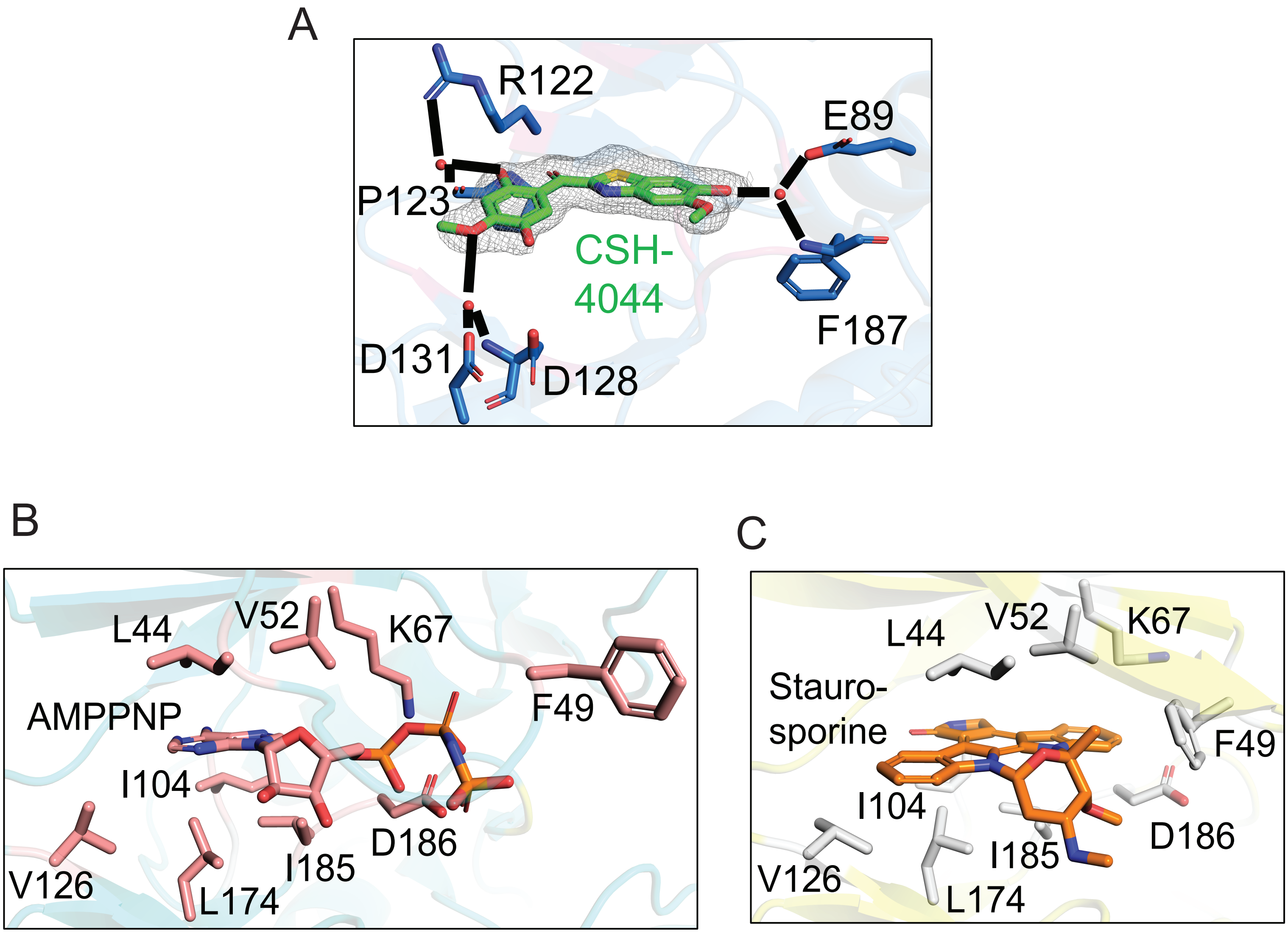


**Figure S2** (A) Close-up view of the water-mediated interaction network in CSH-4044 binding in PIM1. Zoom-in view of (B) AMP-PNP and (C) Staurosporine inhibitor bound to PIM1 kinase. The hydrophobic residues stabilizing the AMP-PNP nucleotide and staurosporine are shown as sticks. Phe49 is repositioned in the staurosporine complex compared to the structure with AMP-PNP, to close the pocket. This repositioning is also observed in the complex with CSH-4044.

### Supplementary Fig 3

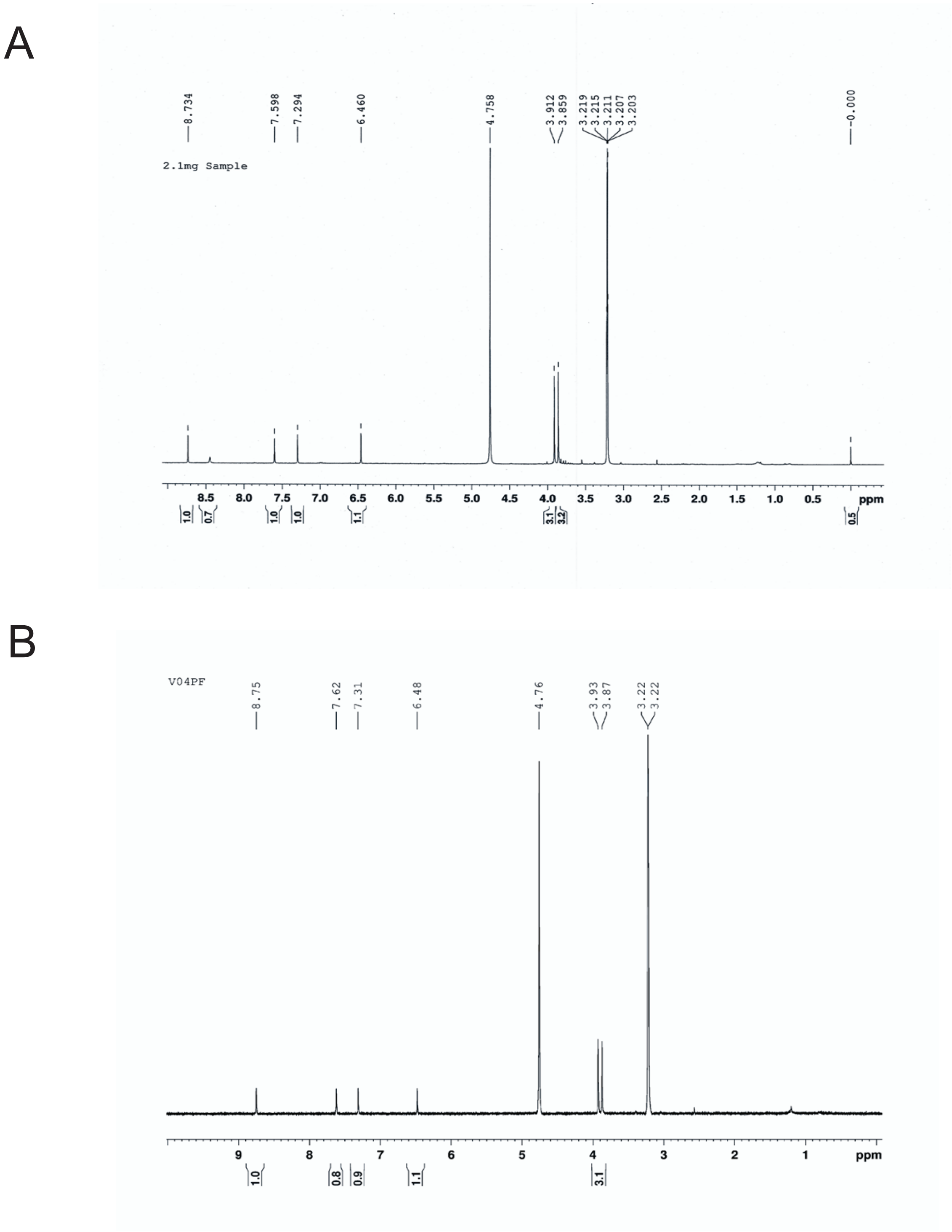


**Figure S3**

**H-**NMR spectrum of the isolated (A) and synthesized (B) inhibitor molecules.

Supplementary Fig 4

A

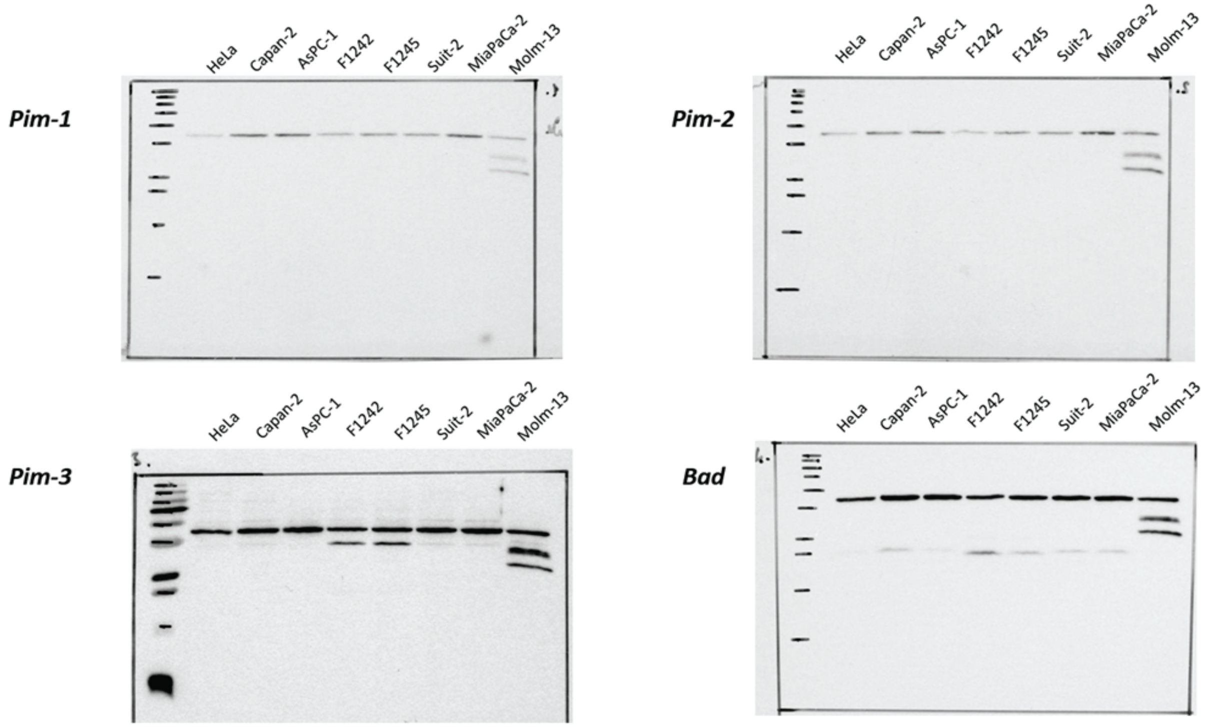


**Figure S4**

**(A)** Immunoblot screening of PIM1, PIM2, PIM3 and BAD in different cell lines
